# Evaluating rapid extraction methods for recovering ancient DNA from archaeological sediments

**DOI:** 10.1101/2025.11.13.688359

**Authors:** Nihan Dilşad Dağtaş, Azhar Jabareen, Krzysztof Cyrek, Maciej T. Krajcarz, Magdalena Krajcarz, Manuel Ángel Rojo-Guerra, Policarpo Sánchez-Yustos, Magdalena Sudoł-Procyk, Shmuel Safra, Viviane Slon

**Affiliations:** The Gray Faculty of Medical & Health Sciences, Tel Aviv University, Israel; The Dan David Center for Human Evolution and Biohistory Research, Tel Aviv University, Israel; Institute of Archaeology, Nicolaus Copernicus University, Toruń, Poland; Institute of Geological Sciences, Polish Academy of Sciences, Warszawa, Poland; Área de Prehistoria, Departamento de Prehistoria, Arqueología, Antropología Social y Ciencias y Técnicas Historiográficas, University of Valladolid, Spain; Faculty of Exact Sciences, School of Mathematical Sciences, Tel Aviv University, Israel

## Abstract

Sedimentary ancient DNA (aDNA) provides a powerful means to reconstruct past environments and populations. However, sedimentary aDNA research is hindered by low success rates due to poor DNA preservation, the largely opportunistic nature of field sampling, and long delays between sampling and results—meaning that substantial time and effort are often spent on samples that eventually yield little or no aDNA. On-site screening methods could thus be particularly valuable when site access is temporary (e.g., seasonal excavations, salvage operations, or in regions affected by geopolitical instability), where delays or poor preservation can mean missed opportunities to recover optimal samples. Here, we compared three rapid DNA extraction protocols—an Amicon ultrafiltration method, an alkaline lysis approach, and a cellulose paper-binding method—across three sediment types (clay, sand, and soil), benchmarking them against a silica-based protocol optimized for ancient sediments but designed for laboratory rather than field conditions. We found that the Amicon-based method, originally validated on ancient bones and coprolites, and which we modified here to enhance the recovery of very short fragments, recovered from sediments both short synthetic DNA and modern DNA sheared to mimic aDNA, outperforming the other rapid methods. When applied to Pleistocene and Holocene archaeological sediments, this method successfully recovered amplifiable aDNA within approximately three hours. While not a replacement for laboratory extractions, this rapid, portable, non-hazardous approach adds a practical tool to the sedimentary aDNA research toolkit, enabling rapid on-site assessment of aDNA preservation and supporting informed sampling in time-sensitive or logistically constrained excavations.

## 1. Introduction

Ancient DNA (aDNA) can offer a unique window into the past, revealing evolutionary processes in ways that modern DNA cannot. However, the absence of active DNA repair mechanisms after death allows damage to accumulate over time (Dabney *et al*., 2013b), and taphonomic processes or storage conditions (Eriksen *et al*., 2025) may severely complicate, and sometimes even prevent, the recovery of aDNA from samples. To overcome this, specialized extraction protocols tailored to aDNA have been developed, and further optimized for different sample types (e.g., Dabney *et al*., 2013a; Rohland *et al*., 2018; Latorre *et al*., 2020; Thomsen *et al*., 2009; Oskam & Bunce, 2012). These DNA extraction protocols are generally composed of two steps: incubating a subsample in lysis buffer (typically overnight) to release DNA into the solution and purifying it from other cellular components and contaminants. Depending on the specific protocol used, the entire process can take up to a few days.

One of the sample types increasingly employed in aDNA research is sediments. Compared to the more commonly used skeletal remains, working with sediment samples offers several advantages. These include being independent of rare fossil material and causing minimal disturbance to archaeological remains, involving only small-scale sampling of sediment layers; the ability to investigate broader temporal scales compared with studies limited to a few fossils; the ubiquity of sediments at excavation sites; and the potential to recover genetic material from a wider range of organisms within a single sample. Thus, the past few years have seen a growing number of landmark studies employing sediment samples to study the past. For instance, genetic analyses on sediments revealed a previously unknown population turnover and genetic diversity decline among Neanderthals at Galería de las Estatuas, Spain (Vernot *et al*., 2021). Similarly, the systematic sampling of hundreds of sediment specimens from Denisova Cave demonstrated that Denisovans, Neanderthals, and anatomically modern humans occupied the site for millennia— sometimes overlapping in time, sometimes replacing one another—and that comparable population shifts also occurred in bears and hyenas (Zavala *et al*., 2021). More recently, the recovery of the oldest DNA obtained to date (Kjær *et al*., 2022) enabled the reconstruction of the flora and fauna inhabiting Greenland two million years ago.

Despite these advantages, sediment samples also present notable challenges compared with fossil remains. Ancient DNA preserved in sediments is typically present at very low concentrations, highly fragmented, can be subject to vertical movement across layers, and is often co-extracted with environmental inhibitors (Pedersen *et al*., 2015). Consequently, the application of robust extraction protocols is essential for recovering usable genetic information from sediments. A handful of such protocols (Rohland *et al*., 2018, Epp *et al*., 2019, Murchie *et al*., 2020, Wang *et al*., 2021) have been successfully used in multiple studies (e.g., Curtin *et al*., 2021, Massilani *et al*., 2022, Buchwald *et al*., 2024, Murchie *et al*., 2021, Zampirolo *et al*., 2024), however, they take approximately 16-24 hours to complete, while requiring heavy laboratory equipment such as high-speed centrifuges and temperature-controlled incubators.

Field-applicable extraction methods offer several strategic advantages (e.g. Utge *et al*., 2020, Selz *et al*., 2023, Kozaczek *et al*., 2025). Rapid on-site processing can provide an immediate initial assessment of DNA preservation, enabling targeted sampling of layers or micro-locations with the highest potential. This improves efficiency, avoids processing poorly preserved samples, and shortens the time from excavation to analysis. Indeed, many sites are located in remote areas without cold-chain infrastructure, and transporting unprocessed sediment to distant laboratories can take days to weeks, increasing the risk of further DNA degradation or contamination. Moreover, waiting for laboratory-based results can delay critical decisions about sampling strategy. On-site methods can be especially valuable in contexts where site access is temporary— for example, during seasonal excavations, under limited excavation permits, in salvage operations, or in regions affected by geopolitical instability—where delays can mean losing the opportunity to recover optimal samples.

To be suitable for field use, a DNA extraction method should ideally be rapid, require only limited and portable equipment, and avoid the use of hazardous reagents (such as guanidine hydrochloride, N-lauroylsarcosine, or phenol) that are only safe to use in laboratories equipped with fume hoods. At the same time, an ideal field-based extraction method should remain sufficiently efficient in recovering DNA with the typical characteristics of aDNA - low abundance, short fragment lengths, and chemical damage. Here, we compared three types of rapid DNA extraction protocols (an Amicon ultrafiltration method, an alkaline lysis method, and a cellulose paper-based binding method: Utge *et al*., 2020; Truett *et al*., 2000; Mason & Botella, 2020, respectively) on three sediment matrix types (clay, sand and soil) to determine which method yields the most amount of DNA, based on both DNA concentration and fragment lengths. Note that of the three rapid protocols tested, only one (Utge *et al*., 2020) was previously tested on ancient samples; and that none were developed nor optimized for ancient sediments. We compared their performance to that of the Rohland *et al*. (2018) extraction method, which was tested and optimized for ancient sediments, but is not readily applicable to field settings. By directly comparing these rapid protocols across different sedimentary matrices, we aimed to identify an extraction method that balances the efficient recovery of authentic aDNA, with the practical constraints of archaeological fieldwork. Lastly, we applied the most promising method— with modifications developed here—to a set of archaeological sediment samples previously shown to contain aDNA, to assess its performance on authentic material.

## 2. Methods

### 2.1 Experimental setup

The performance of three rapid extraction protocols was compared to the Rohland *et al*. (2018) method, which we consider here as the ‘gold-standard’ (henceforth: “Method 1”). Initial evaluation was performed using three ‘mock’ sediment samples - clay, sand, and soil - representing common components of archaeological sediments (Fig. 1). Pure clay (100% red clay; Kibbutz Urim, Israel) and garden soil (standard potting mix) were obtained from commercial sources, and sand was collected from a public beach in Tel Aviv-Yafo, Israel. None of these ‘mock’ samples underwent any laboratory pretreatment. All experiments were performed in the Ancient DNA facilities at the Dan David Center for Human Evolution and Biohistory Research at Tel Aviv University, Israel.

**Figure 1.**
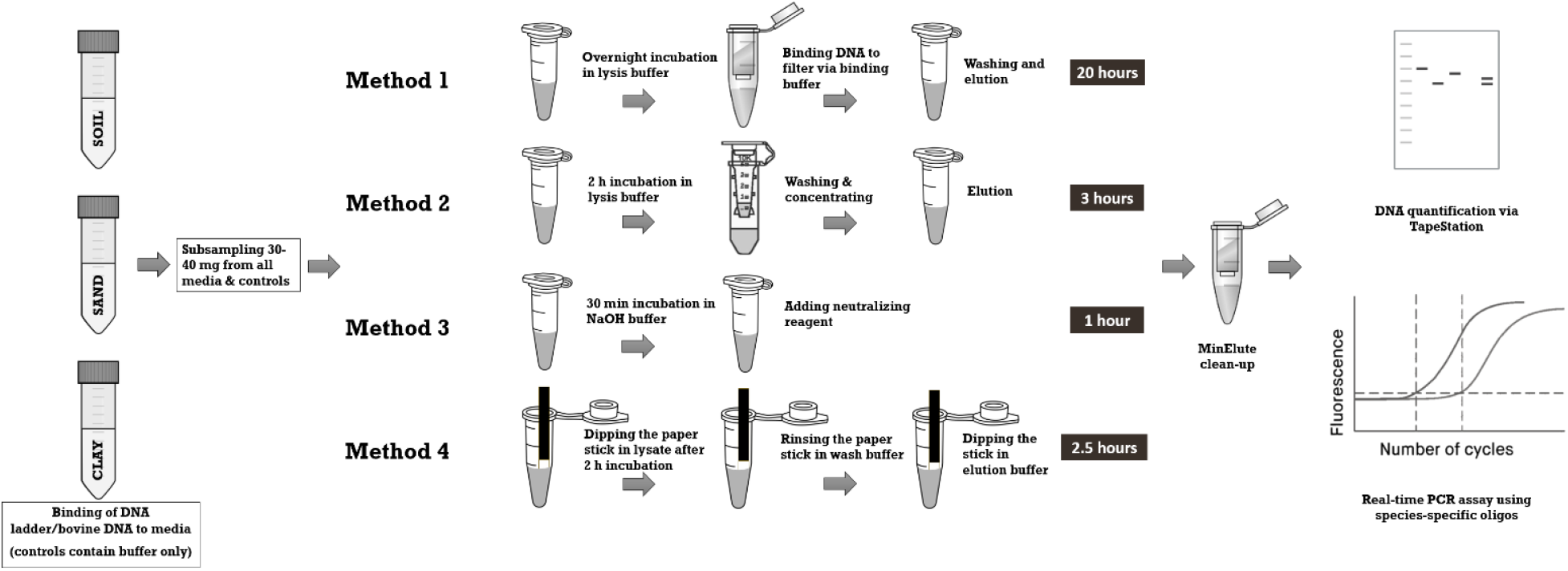
Schematic overview of the experimental setup. In Experiment 1, an artificial Ultra Low Range DNA ladder was bound to the three media types; while in Experiment 2, sheared bovine DNA was bound instead.

### 2.2 DNA binding

Experiment 1: Artificial DNA - To mimic aDNA fragments in size, Ultra Low Range (ULR) DNA ladder (Invitrogen, USA cat. no. 10597012; fragment size range 10-300 bp) was used as the input for extractions (Fig. 1). Forty ug ULR DNA ladder (80 ul) was mixed with 9920 ul of Ringer solution (modified from Slon *et al*., 2017 with a final concentration of 144.3 mM Na^+^, 4.2 mM K^+^, 2.4 mM Ca^+2^, 126.5 mM Cl^−^, 1 mM Mg^+2^, 27 mM Acetate^−^; final volume 10 ml) and added to 1.5 g of each medium - clay, sand and soil. As a negative (no DNA) control, 3.33 ml of Ringer solution was added to 0.5 g of each medium. To allow for sufficient time for DNA binding, all tubes were left for an overnight incubation with rotation at room temperature. Incubation was stopped after ∼22 hours and all tubes were centrifuged at 3220 g for 4 minutes. Supernatants were transferred to clean tubes, and all pellets were washed three times with EBT buffer (10 mM Tris-HCl pH 8.0, 0.05% Tween 20). In every wash step, 10 ml and 3.33 ml of EBT buffer was added to the pellets (for media and media controls, respectively), vortexed and incubated for 5 minutes. Then, the tubes were centrifuged at 3220 g for 1 min, and the supernatants were transferred to clean tubes and kept at −20°C. Between 30-40 mg of the pellets were then subsampled into new 2 ml Lobind tubes (Eppendorf, Germany), and the subsamples were kept at 4°C until used in the extraction tests.

Experiment 2: Bovine DNA - Bovine genomic DNA (EMD Millipore, USA) was sonicated for 1 hour with 175 Watts peak power, 20% duty factor, and 600 cycles/burst settings on Covaris S220 ultrasonicator (Covaris, USA) and bound to the three types of media, as above.

### 2.3 DNA extraction

Following the binding of DNA to all media, three rapid DNA extraction methods were performed in two technical replicates (Utge *et al*., 2020; Truett *et al*., 2000; Mason & Botella, 2020), alongside the Rohland *et al*. (2018) method (see Fig. 1). In short:

Method 1 – Silica-based (Rohland *et al*., 2018) – Overnight incubation of samples at 37°C in lysis buffer (0.45 M EDTA (pH 8), 0.05% (v/v) Tween 20, 0.25 mg/ml proteinase K), followed by binding of DNA to silica spin columns facilitated by a binding buffer (5 M GuHCl, 40% (v/v) isopropanol, 0.12 M sodium acetate, 0.05% (v/v) Tween 20). After two washing steps using 750 ul wash buffer (Buffer PE, Qiagen, Germany), elution in 50 ul elution buffer (0.01 M Tris-HCl (pH 8), 0.001 M EDTA (pH 8), 0.05% (v/v) Tween 20).

Method 2 – Amicon-based (modified from Utge *et al*., 2020) – Two-hour incubation of samples at 42°C in lysis buffer (0.25 M EDTA, 10 mM Tris-EDTA (pH 8), 0.2% SDS [instead of N-lauryl-sarcosyl in the original protocol], 200 ug/ml proteinase K), followed by purification of DNA in Amicon ultra-0.5 ml 10 kDa centrifugal filter units [instead of 30 kDa, see Supplementary Figure S1, Supplementary Table S1] by rinsing 3 times [instead of 4 times as in the original protocol] with ultrapure water. DNA was later recovered in ∼40 ul by centrifugation of the filter unit in reverse position.

Method 3 – HotSHOT: Hot sodium hydroxide (Truett *et al*., 2000) – Half an hour incubation of samples at 95°C in alkaline lysis reagent (25 mM NaOH, 0.2 mM disodium EDTA, pH 12). Then a cooling of samples to 4°C, followed by the addition of a neutralizing reagent (40 mM Tris-HCl, pH 5). The final volume of the extract was approx. 150 ul.

Method 4 – Cellulose paper-based (Mason & Botella, 2020) – After adding lysis buffer (recipe as in Rohland *et al*., 2018) to samples, tubes were vortexed and incubated with rotation for 2 h at room temperature. This was to ensure the release of DNA fragments into the solution, although the original study suggests a lysis step of max. of 30 min. Tubes were centrifuged for 30 s at 13000 rpm. Custom-made cellulose-paper sticks (0.5 x 7-8 cm) were then dipped in the lysates— until the DNA binding zone was submerged—for ∼5 s. Then, the sticks were dipped in 800 ul wash buffer (10 mM Tris (pH 8)) 5 times for a total of ∼5 s, followed by a dip in 200 ul elution buffer (recipe as in (Rohland *et al*., 2018)) for ∼10 s.

All supernatants and extracts were later cleaned up using MinElute PCR Purification kit (Qiagen, Germany cat. no. 28004) following the manufacturer’s protocol and eluted in 30 ul Qiagen EB buffer. All extraction batches included a positive (ULR DNA ladder) and a negative control (dH_2_O).

### 2.4 DNA quantification and analysis

To confirm that the DNA was successfully bound to the media and to assess the performance of the various extraction methods, all clean extracts were run on TapeStation using a D1000 screentape (Agilent, USA cat. no. 5067-5582). DNA concentrations and fragment sizes were measured via the TapeStation Analysis software (v. 5.1). We chose a fragment size range of 35-350 bp for comparison among protocols to exclude the lower marker of the D1000 screentape (25 bp), and to allow for slight measurement shifts that may occur in different TapeStation runs.

In addition, to test the amplifiability of the extracted DNA fragments recovered using different protocols, a real-time PCR assay was performed on the bovine DNA bound to the different media types (Experiment 2). To optimize the assay, we tested between 0.5 and 2 ul of DNA input. Besides the DNA input, the assay contained 10 ul 2X Fast SYBR Green Master Mix (Thermo Fisher Scientific, USA), 900 nM of forward and reverse primers, and 250 nM of TaqMan probe in a 20 ul reaction volume and ran on Azure Cielo Real-Time PCR (Azure Biosystems, USA). Two sets of primers and TaqMan MGB probes (labelled with FAM) designed for *Bos primigenius* and *Crocuta crocuta* (Utge *et al*., 2020) were used, the latter chosen as a negative control as no amplification was expected when using bovine DNA as input for the assay. Molecular grade water was used as qPCR blank and sheared bovine DNA (0.5 ul/reaction) was used as qPCR positive control. Reaction conditions were as follows (from Utge *et al*., 2020): 2 min at 50°C, 20 s at 95°C, then 40 cycles of (3 s at 95°C, 30 s at 60°C). For Method 4 after elution, we also dipped each paper in 40 ul qPCR master mix and incubated for 30 min to possibly release any leftover DNA attached to papers and included these mixes in the qPCR run (Fig. 4D).

We tested whether there is any significant difference between the DNA yields obtained via the four extraction protocols using a pairwise Wilcoxon test, implemented in the rstatix package in RStudio version 2024.12.0+467.

### 2.5 Application to archaeological sediment samples

Based on the results of these tests, nine sediment samples from three archaeological sites were subjected to DNA extraction using Method 2 (Utge *et al*. 2020 protocol, with the modifications mentioned in section 2.3). These included four samples from Perspektywiczna Cave (Poland); three samples from Cueva Millán (Spain); and two samples from Biśnik Cave (Poland) (Table 1). These samples were chosen because prior screening (unpublished data) indicated the presence of preserved aDNA when testing for mammalian mitochondrial DNA (mtDNA; following the strategy described in Slon *et al*., 2017). The selection spans contexts from the Middle Pleistocene to the Holocene, includes both caves and a rock shelter, and covers distinct geographical regions in Europe.

**Table 1.** Archaeological context for the sediment samples tested.

| Sample ID | Context | Site | Chronostratigraphy | Type of site | Main ref.s for the sites/layers | Oligos expected to amplify, based on previous seq. data |
| --- | --- | --- | --- | --- | --- | --- |
| S584 | Layer 8 | Perspektywiczna Cave | Lower Holocene | Cave (near the entrance) | Krajcarz *et al.* 2023 | Cervus |
| S603 | Layer 5 |  | Holocene? | Cave (near the entrance) | Krajcarz *et al.* 2023 | Crocutea |
| S612 | Layer 7c |  | Upper Pleistocene | Cave | Krajcarz *et al.* 2023 | Crocutea |
| S613 | Layer 7c |  | Upper Pleistocene | Cave | Krajcarz *et al.* 2023 | Crocutea |
| S1201 | Layer 1 | Cueva Millán | Upper Pleistocene | Rock shelter | Sánchez-Yustos *et al.* 2024 | Cervus, Crocutea |
| S1202 | Layer 1 |  | Upper Pleistocene | Rock shelter | Sánchez-Yustos *et al.* 2024 | Cervus, Crocutea |
| S1203 | Layer 1 |  | Upper Pleistocene | Rock shelter | Sánchez-Yustos *et al.* 2024 | Cervus, Crocutea |
| S1206 | Layer 15 | Biśnik Cave | Middle Pleistocene | Cave | Krajcarz *et al.* 2014 | None |
| S1207 | Layer 18 |  | Middle Pleistocene | Cave | Krajcarz *et al.* 2014 | None |

Real-time PCR assays were performed as in Section 2.4 (except with only 1 ul DNA input) using *C. crocuta* and/or *Cervus elaphus* primers and TaqMan probes (Utge *et al*., 2020), depending on the taxa previously identified in each sample (Table 1). These oligos (i.e., artificially produced single-stranded DNA fragments used as primers and probes) amplify a 68 bp region on the COX3 gene and a 66 bp region on the CYTB gene, respectively, and both are located on the mitochondrial genome. Note that the two samples from Biśnik Cave, which only yielded ancient Ursidae DNA in a previous sequencing effort, served as negative controls, as no amplification was expected using the two above-mentioned species-specific oligos.

## 3. Results

To confirm that the ULR DNA ladder and the sheared bovine DNA were successfully bound to the three different types of media, supernatants collected after DNA binding as well as wash eluates were cleaned up using MinElute PCR Purification kit, and run on TapeStation using the D1000 screentape. Figure 2 shows that for most cases no DNA is detected in the supernatants, proving that DNA had successfully bound to clay, sand and soil. This provided the basis to perform various DNA extraction protocols on these ‘mock’ sediment samples.

**Figure 2.**
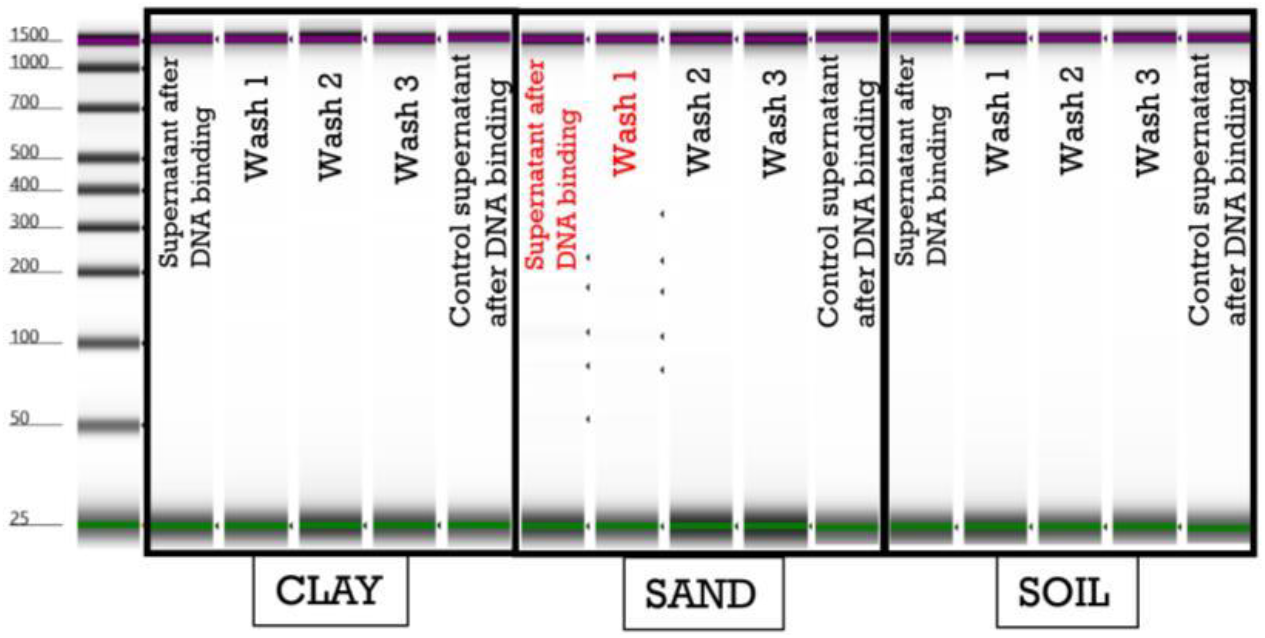
Confirmation of successful binding of DNA to clay, sand and soil. Artificial DNA in the form of an ULR DNA ladder was bound to three types of media. Supernatants collected after the binding step and after each wash step were run on TapeStation. In most cases, no DNA was detected in the supernatants, confirming the successful binding of DNA to the media. Note the rather long but faint bands of artificial DNA in the sand supernatant after binding and in the first wash (highlighted in red), which may be due to the lower DNA binding capacity of sand compared to clay, as seen in Slon *et al*., 2017. None of the negative controls contain background DNA. Note that this is a composite image of two TapeStation runs (images were normalized for exposure and combined digitally for presentation; no alteration of data was performed).

We then extracted DNA from the ‘mock’ sediment samples from Experiment 1 (bound artificial DNA) using the four protocols. As expected (Slon *et al*., 2017; Rohland *et al*., 2018), Method 1 (silica-based) successfully recovered DNA fragments, even of short sizes, from clay and sand (Fig. 3). The apparent lower DNA yields (reflected by fainter bands in the TapeStation run) for sand compared to clay may be due to a weaker initial DNA binding performance (Fig. 2). Extraction of DNA with Method 1 from the three media yielded higher amounts of DNA than with any other tested method (Table 2), although this difference was not statistically significant (pairwise Wilcoxon test, p = 0.25). Apart from a few faint bands, no traces of DNA were discernible in soil extracts, possibly due to inhibitory substances present in commercial garden soil. With Method 2, DNA fragments as short as 35 bp were recovered from clay as well as from sand (Fig. 3). Yields of DNA from soil (5.58–6.48 ng/g) were similar to the control (4.9 ng/g), showing no consistent evidence of recovery above background (Table 2). A poorer performance was noted for Method 3, which showed no detectable DNA from clay or soil, and yielded only few visible DNA fragments from sand (Fig. 3). Method 4 did not produce any detectable DNA above the background from clay or sand (Fig. 3), and only very low yields of DNA from soil (Table 2). It took approximately 20, 3, 1 and 2.5 hours to complete each protocol, respectively.

**Table 2.** Summary statistics of DNA recovery for all tested extraction methods. The DNA concentration is expressed as yield (ng/g) after correcting for subsample amount, and the results are for the fragment size range of 35-350 bp. Replicates 1-2 correspond to two technical replicates, and Control corresponds to the background values measured in the negative controls for each medium. Note that the outlier ‘Replicate 2 in Soil/Method4’ (in bold) was excluded from the statistical analysis due to a faulty TapeStation result.

|  | Clay |  |  |  |  |  |  |  | Sand |  |  |  |  |  |  |  | Soil |  |  |  |  |  |  |  |
| --- | --- | --- | --- | --- | --- | --- | --- | --- | --- | --- | --- | --- | --- | --- | --- | --- | --- | --- | --- | --- | --- | --- | --- | --- |
|  | Method 1 |  | Method 2 |  | Method 3 |  | Method 4 |  | Method 1 |  | Method 2 |  | Method 3 |  | Method 4 |  | Method 1 |  | Method 2 |  | Method 3 |  | Method 4 |  |
|  | Yield (ng/g) | Average size (bp) | Yield (ng/g) | Average size (bp) | Yield (ng/g) | Average size (bp) | Yield (ng/g) | Average size (bp) | Yield (ng/g) | Average size (bp) | Yield (ng/g) | Average size (bp) | Yield (ng/g) | Average size (bp) | Yield (ng/g) | Average size (bp) | Yield (ng/g) | Average size (bp) | Yield (ng/g) | Average size (bp) | Yield (ng/g) | Average size (bp) | Yield (ng/g) | Average size (bp) |
| Replicate 1 | 88.50 | 119 | 51.75 | 123 | 6.90 | 54 | 8.23 | 50 | 33.67 | 94 | 19.55 | 89 | 9.83 | 69 | 5.03 | 63 | 14.83 | 90 | 6.48 | 61 | 4.45 | 61 | 10.20 | 57 |
| Replicate 2 | 74.25 | 119 | 16.73 | 105 | 4.20 | 51 | 6.27 | 52 | 31.50 | 96 | 12.68 | 65 | 7.53 | 86 | 11.67 | 53 | 14.73 | 95 | 5.58 | 51 | 11.37 | 74 | **28.67** | 60 |
| Control | 5.23 | 69 | 7.03 | 48 | 6.73 | 54 | 7.07 | 50 | 5.97 | 62 | 6.08 | 51 | 3.10 | 63 | 7.03 | 49 | 5.53 | 129 | 4.90 | 65 | 4.18 | 63 | 3.65 | 61 |
| Replicate avr. | 81.38 | 119 | 34.24 | 114 | 5.55 | 53 | 7.25 | 51 | 32.58 | 95 | 16.11 | 77 | 8.68 | 78 | 8.35 | 58 | 14.78 | 93 | 6.03 | 56 | 7.91 | 68 | 10.20 | 57 |

**Figure 3.**
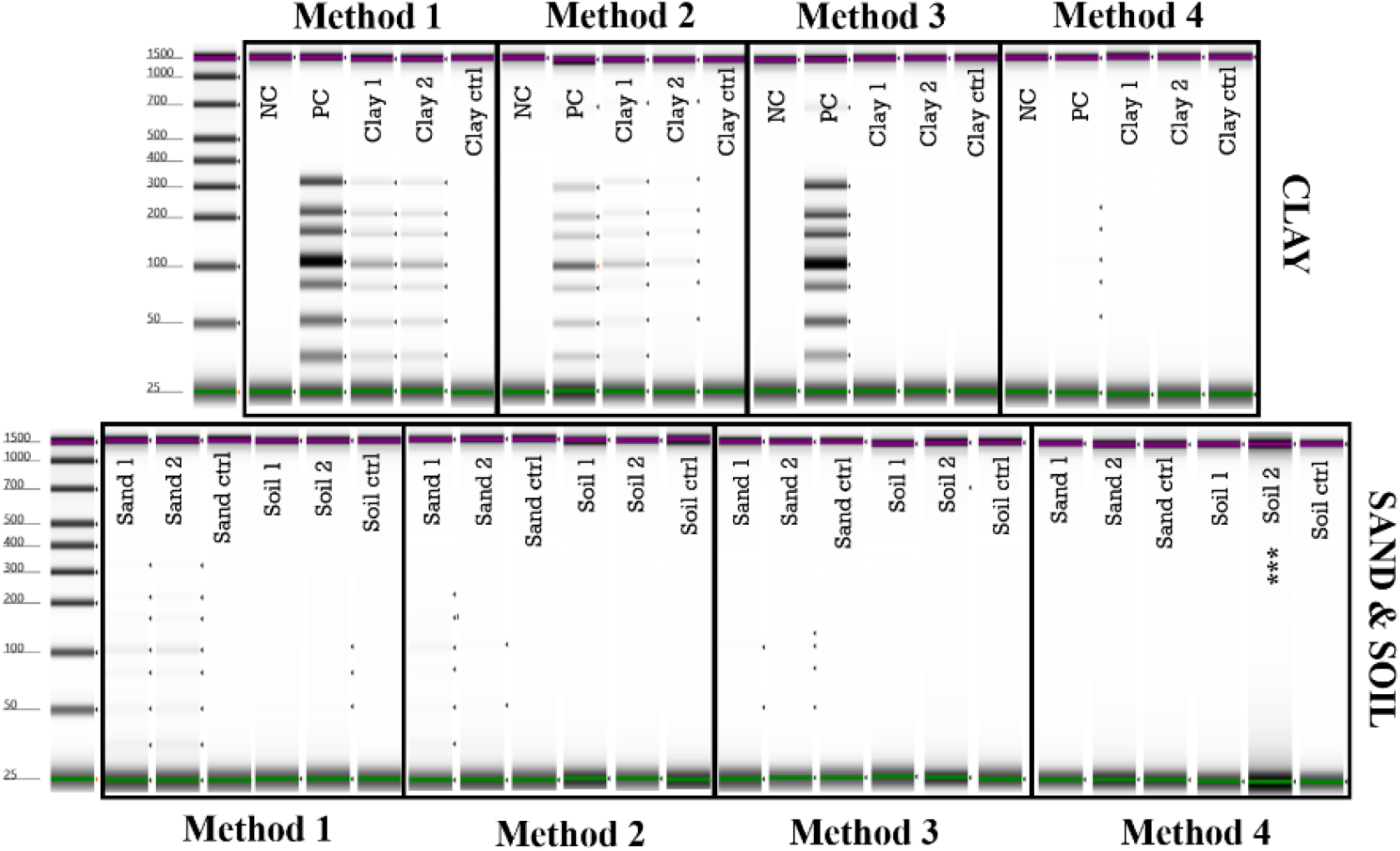
Evaluation of four protocols for extracting DNA from clay, sand and soil. DNA was successfully retrieved from clay (upper half) using Methods 1 and 2, and from sand (lower half) using Methods 1-3. None of the protocols consistently produced detectable DNA from soil (lower half). (NC: negative control; PC: positive control; *** marks the sample excluded from the statistical analysis). Note that this is a composite image of multiple TapeStation runs (images were normalized for exposure and combined digitally for presentation; no alteration of data was performed).

Similar results were found for the second experimental setup, in which sheared bovine DNA was bound to the three types of sedimentary matrices (Figure S2, Table S2; only clay is shown). Based on these results, we performed follow-up experiments with archaeological sediment samples focusing on Method 2 (Utge *et al*., 2020, with modifications).

Electrophoretic readouts (i.e., TapeStation results) can confirm the recovery of detectable DNA and provide approximate fragment-size distributions, but they have limited sensitivity at low copy numbers, are biased against very short fragments, and cannot readily distinguish poor recovery from co-extracted inhibitors. To determine whether the extracts produced with the different protocols are suitable for downstream analyses, we next tested the amplifiability of the recovered DNA fragments by real-time PCR. With the second experimental setup, in which bovine DNA was bound to the three sedimentary matrices, real-time PCR assays detected amplifiable target DNA using oligos designed to amplify aurochs (*B. primigenius)* DNA for all tested conditions and media, but not in any of the negative controls (Fig. 4A-D, Table S3). This amplification was achieved even for the soil medium with Methods 1-3, where the concentration of extracted DNA was shown to be low relative to the clay and sand media (Table 2). Observing the expected 1 cycle difference between the 0.5, 1 and 2 ul DNA inputs used in the reactions supported the efficiency of the qPCR assay (except for Soil/Method4, Table S3). We note that the aurochs TaqMan probe is not as species-specific as previously thought (Utge *et al*., 2020), as it was able to amplify bovine (modern cattle) DNA. Lastly, as expected, when using oligos designed to amplify hyena (*C. crocuta*) DNA, which served as a negative control to verify assay specificity, none of the samples displayed amplifiable DNA. Indeed, none of the positive Cq values for *C. crocuta* indicate true amplification, but rather appear to be noise, as evidenced by the flat, non-amplifying curves in Figure 4A-D.

**Figure 4.**
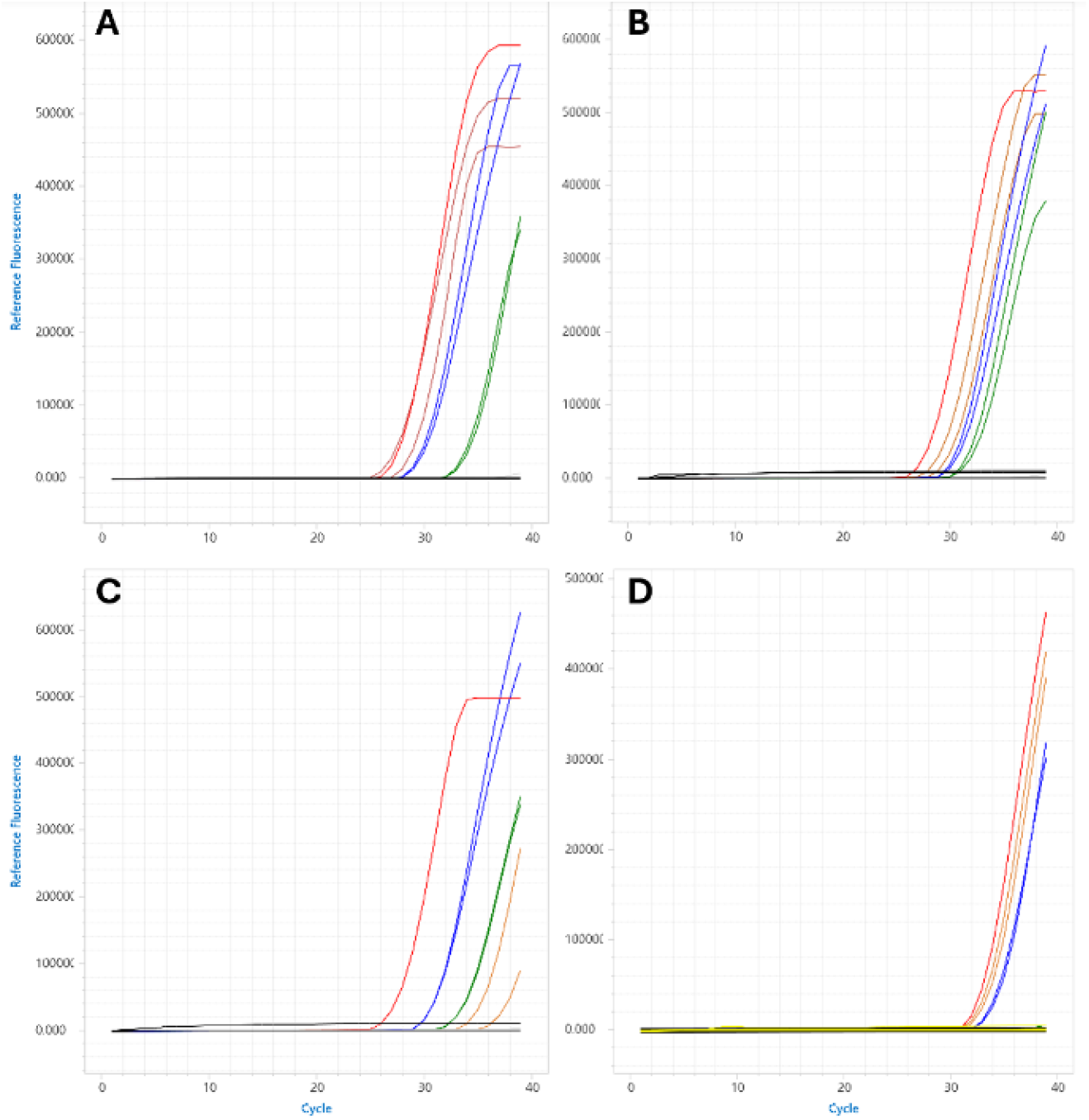
Quantification of bovine DNA extracts via real-time PCR. Bovine DNA was extracted following four different protocols and quantified using *B. primigenius* and *C. crocuta* oligos (the latter used as a negative control). A) Method 1, B) Method 2, C) Method 3, D) Method 4. Amplification curves indicate successful recovery of amplifiable bovine DNA from all tested sediment media (orange: clay replicates; blue: sand replicates; green: soil replicates; red: positive control). Note the straight gray and black lines at Fluorescence 0 (zero) showing no amplification in the negative controls (media controls, extraction negative controls and qPCR blank), nor with the *C. crocuta* oligos, respectively. In D, straight yellow lines indicate no amplification in qPCR master mix tubes into which papers were dipped to release any leftover DNA. Only reactions with 1 ul DNA are shown for clarity).

Lastly, we tested the applicability of Method 2 to a set of nine archaeological sediment samples, amplifying them with *C. crocuta* and/or *C. elaphus* oligos based on the expected presence of relevant ancient taxa in the samples (Table 1). All samples from Perspektywiczna Cave expected to contain ancient Hyaenidae mtDNA (S603, S612, S613) yielded consistent positive results for both replicates with the *C. crocuta* oligos, whereas one sample (S1203) out of three from Cueva Millán displayed an amplification, albeit not reproducible in the second—more diluted—replicate (Fig. 5A, Table S4). Sample S1206, chosen as negative control as no ancient Hyaenidae mtDNA was expected in it (Table 1), showed no amplification; neither did any of the negative controls (Fig. 5A, Table S4). With the *C. elaphus* oligos, we observed late amplification signals in three of the four samples where ancient Cervidae mtDNA is expected (Fig. 5B; Table S4), suggesting potential target DNA. However, these signals were weak, appeared at high Cq values, and were not consistently reproducible across replicates. To assess PCR inhibition in Cueva Millán samples, second replicates (S1201-S1203) were run using a diluted template, but these did not yield stronger signals. Sample S584, from a different site, produced only a single late, non-reproducible amplification. As with the *C. crocuta* assay, neither sample S1207 which served as a negative control (Table 1), nor any of the negative controls, displayed an amplification (Fig. 5B; Table S4). Taken together, these results suggest that the issue with the *C. elaphus* assay is more likely related to assay optimization than to inhibition, and that further refinement of qPCR conditions may improve detection. In particular, we note that the use of SYBR Green-based master mixes in TaqMan assays was likely sub-optimal and may have reduced amplification efficiency (Cheah *et al*., 2010). Future assays employing the TaqMan master mix may yield stronger and more consistent signals.

**Figure 5.**
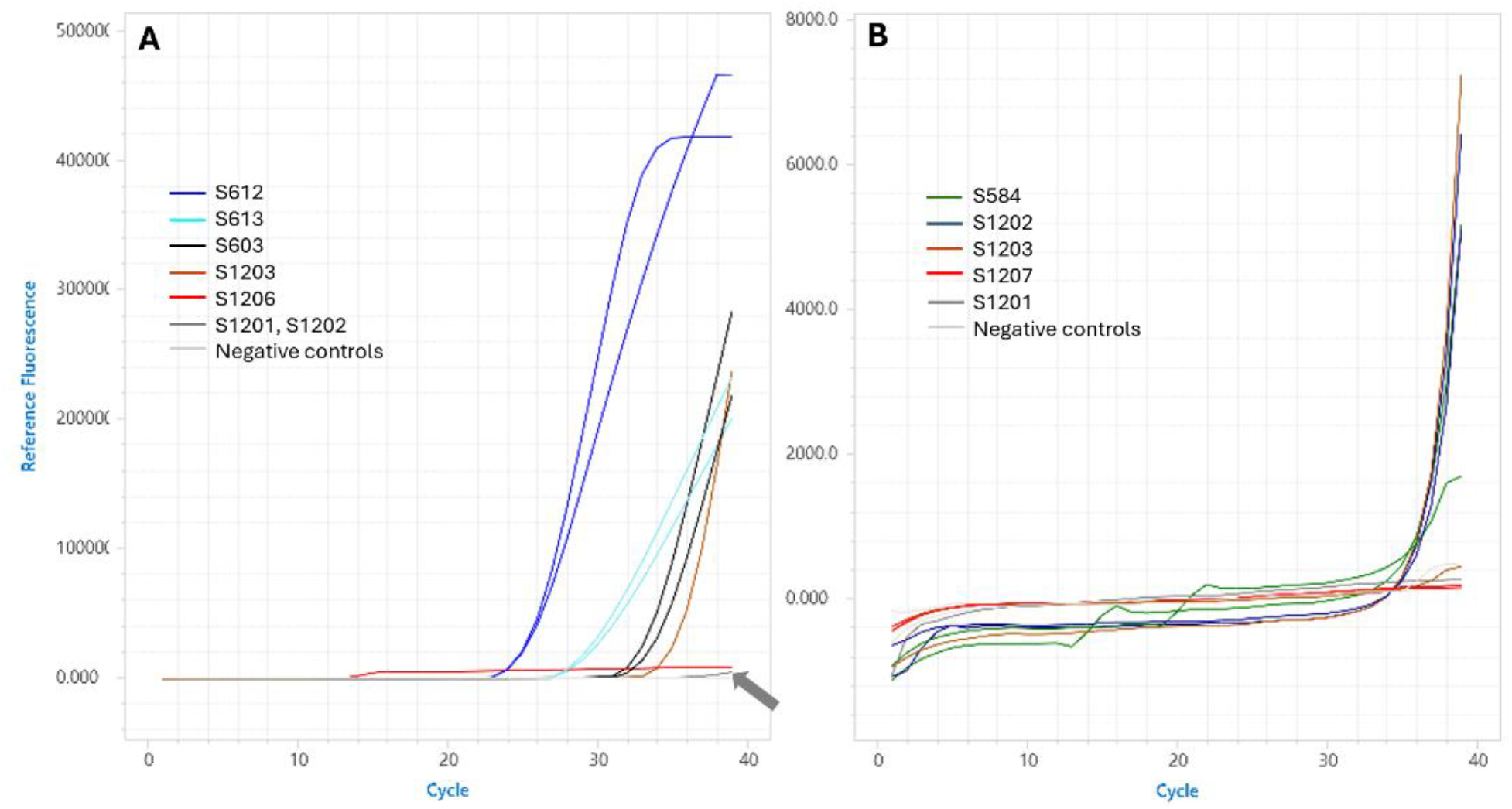
Quantification of archaeological sediments via real-time PCR. A) Amplification using *C. crocuta* oligos. An amplification was obtained for four samples (S603 in black, S612 in dark blue, S613 in light blue, S1203 in orange) where ancient Hyaenidae mtDNA was expected (Table 1). Samples that did not amplify (S1201, S1202) are in dark gray. The arrow shows the dark gray line of S1201-Replicate2, a late, non-reproducible signal with a Cq value of 39 (Table S4), which is interpreted as background noise possibly caused by probe degradation towards the end of the run. B) Amplification using *C. elaphus* oligos. A late amplification signal was observed for three samples (S584 in green, S1202 in dark blue, S1203 in orange) where ancient Cervidae mtDNA was expected (Table 1). The sample that did not amplify (S1201) is marked in dark gray. Note the straight light gray and red lines at Fluorescence 0 (zero) showing no amplification in the negative controls, nor in the samples expected to lack the target taxon, respectively. Legends show sample identifiers with their corresponding colors.

In summary, real-time PCR results performed on DNA extracts from archaeological sediment samples were in line with expectations based on sequencing data, in that samples with sequenced DNA fragments pertaining to a particular taxon were also positive in the relevant quantitative assay in several cases. These results demonstrate that Method 2 can successfully recover amplifiable sedimentary aDNA from archaeological contexts and is suitable for application under field conditions.

## 4. Discussion

This study compared the DNA recovery performance of four extraction methods on mock and archaeological sediment samples to evaluate their suitability for field application. Our results demonstrate that the protocol developed by Utge *et al*. (2020) for bones and coprolites is also applicable to archaeological sediment samples, with the modifications developed here (section 2.3, Table S1, Figure S1). Across different sediment types, this method produced DNA yields and fragment sizes that, while generally lower than those obtained with the laboratory-optimized Rohland *et al*. (2018) protocol, outperformed other rapid methods tested. We also showed that the method is compatible with downstream amplification of aDNA fragments recovered from sediment samples, as shown using bovine DNA sheared to mimic the short fragment lengths typical of aDNA.

In addition, when applied to nine archaeological sediment samples from Poland and Spain, spanning the Middle Pleistocene to the Holocene, this rapid extraction method produced extracts suitable for downstream amplification. The positive amplifications obtained with the *C. crocuta* oligos are consistent with the presence of ancient Hyaenidae mtDNA in the sediments, as well as the presence of *C. crocuta* bones and teeth at Perspektywiczna Cave (Krajcarz *et al*., 2023). We note that the assay was developed from the reference mitochondrial genome of cave hyena (Late Pleistocene *C. crocuta* from Coumère Cave in France, NCBI accession number NC_020670.1; Bon *et al*., 2012), and while sequence similarity could, in principle, allow amplification of other *Crocuta* lineages, modern contamination can be excluded given that spotted hyenas are now restricted to Africa and disappeared from these parts of Europe between around 40,000-33,000 years ago (Krajcarz *et al*., 2023; Stuart & Lister, 2014). The signals obtained with *C. elaphus* oligos in some of the tested samples likely reflect ancient Cervidae mtDNA, though a modern origin cannot be ruled out based on these data alone, as red deer and related species are still extant in Europe. Overall, these results indicate that the applied rapid extraction method is capable of recovering amplifiable aDNA from Pleistocene archaeological sediments.

Although this method is unlikely to replace widely used DNA extraction protocols such as those by Dabney *et al*. (2013a), Rohland *et al*. (2018) or Murchie *et al*. (2020), it may serve as a practical alternative in contexts where rapid results are required and full laboratory infrastructure is unavailable. Its relatively short processing time, limited equipment needs, and avoidance of hazardous reagents make it particularly suitable for use in mobile laboratories, temporary excavation setups, or other field-based applications. The demonstrated ability to recover amplifiable sedimentary aDNA under these conditions supports its potential role in enabling informed targeted sampling and on-site screening in archaeological and palaeoenvironmental projects. In this way, an on-site rapid DNA extraction approach contributes to the growing toolkit for investigating the genetic record preserved in sediments, reinforcing the central role of sedimentary aDNA in reconstructing the environmental past.

## Acknowledgements

We thank Dheeraj Chaudhary and Valentina Vanghi for providing DNA extracts for an initial test; Aviva Lindell for technical help in the laboratory; Natalia Aleshkevich for logistic support; and Jean-Marc Elalouf for consultation about the primers. This study was funded by the John Templeton Foundation (Grant no. 62571 to S.S. and V.S.). Research and fieldwork at Cueva Millán are coordinated by the research group DURIUS (University of Valladolid) within the framework of the projects Arqueosabinares, funded by the Junta de Castilla y León (Spain), and PIONEERS (Ref. PID2024-162134NB-I00), funded by the Ministerio de Ciencia, Innovación y Universidades (Spain), both led by P.S.Y. and M.A.R.G.

## Author contributions

N.D.D. and V.S. conceived the study and designed the experimental work; N.D.D. carried out the laboratory work, with support from A.J.; N.D.D. analyzed the data, under the supervision of V.S.; K.C., M.T.K., M.K., M.A.R.G., P.S.Y. and M.S.P. provided archaeological samples and contextual information; M.A.R.G., P.S.Y., S.S. and V.S. acquired funding; N.D.D. and V.S. wrote the manuscript with input from all authors.

