## Supplemental figures for "Evaluating rapid extraction methods for recovering ancient DNA from archaeological sediments"

### Supplementary Information

Comparison of Amicon filters used in Method 2

To evaluate the performance of the 30 kDa Amicon filters employed in the original study where Method 2 was developed (Utge *et al*., 2020), with respect to the retention of short DNA fragments, we compared filters with molecular weight cutoffs of 10 kDa against 30 kDa (UFC501024 and UFC503024, respectively, from Merck, Germany). Our objective was to identify the most effective way of recovering DNA fragments as short as 35 bp, which we defined as the lower bound in our analyses. Using the formulas [(no. of nucleotides × 303.7) + 79] for ssDNA, and [(no. of nucleotides × 607.4) + 158] for dsDNA, the molecular weights of 35 bp fragments were estimated to be 10.7 kDa and 21.4 kDa, respectively (formulas taken from Thermo Fisher DNA and RNA Molecular Weights and Conversions tool). Given that Amicon filters retain molecules larger than their nominal cutoff, we inferred that the 10 kDa filters would provide greater efficiency for capturing short fragments—more likely to represent genuinely ancient DNA fragments (Dabney *et al*., 2013).

To do so, 50mg subsamples of the DNA-bound clay medium and a positive control (ULR DNA ladder) were processed, in duplicates, following the protocol described by Utge *et al*., (2020) for each of the two tested filter types. A negative control ($dH_2O$) was carried along. The results demonstrate that filters with a 10 kDa cutoff retained more DNA (as evidenced by the intensity of the DNA bands) from both the clay and the positive control. In particular, the 10 kDa filters are more effective at retaining DNA molecules in the relevant size range, including ones as short as 35 bp (Figure S1, Table S1).

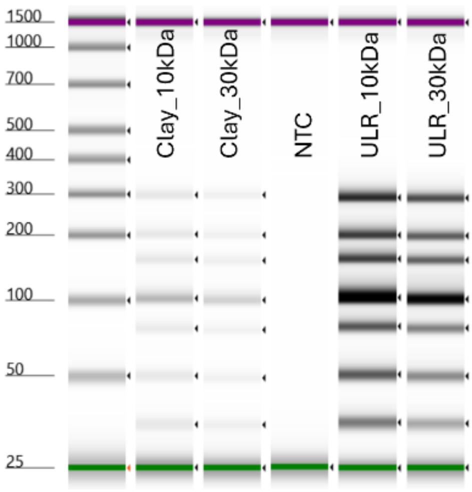


**Figure S1. Comparison of the performance of Method 2 in recovering DNA fragments from clay, using two Amicon filters with different molecular weight cutoffs (10 kDa and 30 kDa).** NTC: negative control; ULR: Ultra Low Range DNA ladder.

Quantification of bovine DNA extracts

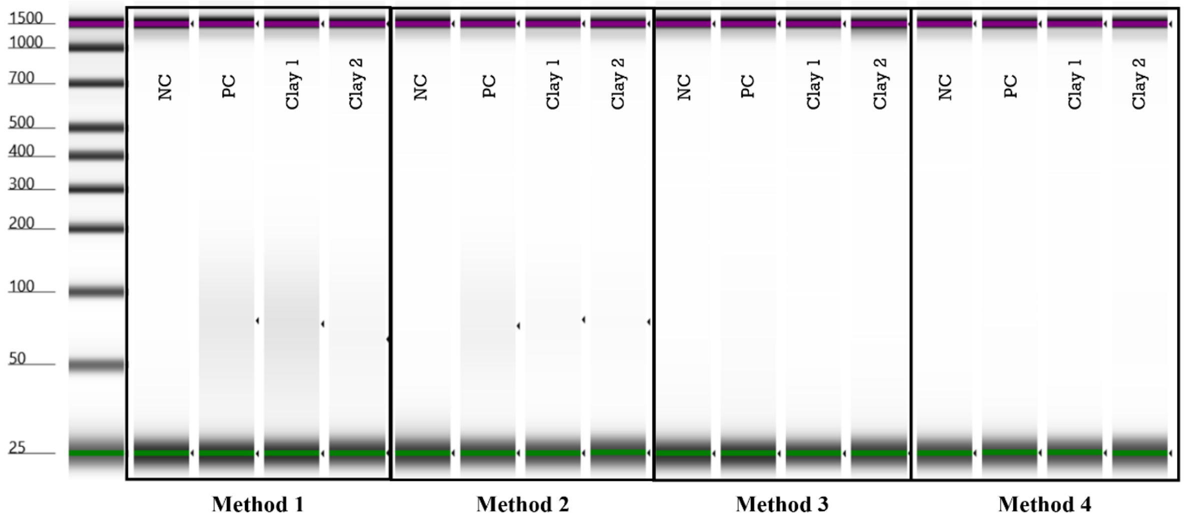


**Figure S2. The TapeStation image of the samples in Table S2.** NC: negative control; PC: positive control; Clay 1 and 2 are technical replicates.
